# Curcumin combined with verapamil improve cardiovascular phenotype of a Williams-Beuren Syndrome mice model reducing oxidative stress

**DOI:** 10.1101/2022.12.19.520405

**Authors:** Noura Abdalla, Paula Ortiz-Romero, Isaac Rodriguez-Rovira, Luis A. Pérez-Jurado, Gustavo Egea, Victoria Campuzano

## Abstract

Williams-Beuren syndrome (WBS) is a rare neurodevelopmental disorder with hallmarks in the cardiovascular manifestations and no well-defined therapeutic strategies. We investigated the progression of cardiovascular phenotype present in CD (complete deletion) mice, a murine model of WBS, after chronic treatment with curcumin and verapamil, both compounds with positive effects on related pathologies. Treatment was administered orally dissolved in drinking water. After measuring “in vivo” systolic blood pressure, aortic and left ventricular myocardium were histological and molecular analysed to determine the effects of treatment and its underlying mechanism in CD mice. We observed upregulated xanthine oxidoreductase (XOR) expression in both aortic and left ventricular myocardium of CD mice. This overexpression is concomitant with increased levels of nitrated proteins as example of byproduct-mediated oxidative stress damage, indicating of XOR-generated oxidative stress impact on the pathophysiology of cardiovascular manifestations in WBS, and suggesting that the inhibition of XOR and/or oxidative stress damage could help to ameliorate the severe cardiovascular injuries presented in WBS.

## Introduction

The cardiovascular pathology present in Williams-Beuren syndrome (WBS, OMIM 194050) is characterized by the presence of generalized arteriopathy associated with hypertrophy of vascular smooth muscle cells (SVMC), disorganized, fragmented elastic fibers, and increased number of lamellar structures (Pober, 2010). Although stenotic lesions are seen in the thoracic aorta, pulmonary artery, and renal artery (Collins, 2018), ascending aortic stenosis (SVAS), which affects more than 70% of patients (Collins, 2018), is the most common cause. most common cardiovascular abnormality. Severe SVAS can lead to cardiac hypertrophy and increases the risk of complications such as stroke and sudden death (Wessel et al., 2004; Collins, 2018).The complete deletion (CD) mouse model of WBS presents a cardiovascular phenotype very similar to that seen in patients with presence of aortic stenosis, hypercardiopathy and hyperthensión (Segura-Puimedon et al., 2014; Jiménez-Altayó et al., 2020). Morever this mouse model has been used to assay potential therapeutic interventions (Ortiz-Romero et al., 2018, 2021).

Oxidative stress is a significant contributing factor to the pathogenesis of several human diseases and considered one of the major cardiovascular risk factors impairing endothelial function, and inducing arterial remodeling and vascular inflammation (Senoner and Dichtl, 2019). Oxidative stress is characterized by the overproduction of reactive oxygen species (ROS), which can damage all components of the cell including proteins, lipids, and DNA (Costa et al., 2021). In WBS patients, ROS levels have been described as strong determinants of hypertension risk and vascular stiffness (Del Campo et al., 2006; Kozel et al., 2014). This hypothesis was supported by observations in other WBS murine model defined by the partial distal deletion (DD) (Campuzano et al., 2012). Of notice, cardiovascular injuries reported in the murine model of complete deletion (CD) are significantly minor compared to those presented in the DD model. This fact has been explained in part by the reduced expression of NCF1 in affected tissues of CD (Segura-Puimedon et al., 2014; Ortiz-Romero et al., 2018). However, both the heart and the aorta of the CD model continue to show high levels of oxidative stress (Ortiz-Romero et al., 2018; Jiménez-Altayó et al., 2020).

Biochemical, molecular and pharmacological studies further implicate xanthine oxidoreductase (XOR) as a source of ROS in the cardiovascular system (Kinugasa et al., 2003; Engberding et al., 2004; Polito et al., 2021). Although the participation of XOR in WBS is unknown, related diseases as Loeys-Dietz and Marfan syndromes showed XOR protein levels increased in aortic samples (Soto et al., 2020; Rodríguez-Rovira et al., 2022). Previous studies have reported that ROS generated by hypoxanthine and xanthine oxidase were inhibited by curcumin (Akter et al., 2020) and the calcium channel blocker verapamil hydrochloride (Liang et al., 2000).

Curcumin (CUR) is a natural yellow pigment, which has attracted much attention in recent years owing to its wide spectrum of biological activities, including antioxidant, anti-inflammatory, anti-tumor, or anti-microbial activities, (Llano et al., 2019; Bozkurt et al., 2022). Besides, CUR improved cardiovascular structure and function, especially with the normalization of systolic blood pressure and collagen deposition in rats with diet-induced metabolic syndrome (du Preez et al., 2019). In addition, CUR has previously shown cardiac protection in front of palmitate and high fat diet mediated the activation of the nuclear factor erythroid 2 (NRF2) (Zeng et al., 2015). On the other hand, calcium channel blockers, such as Verapamil (VER) are used as antihypertensive drugs for long time acting as peripheral vasodilators by blocking entry of calcium through voltage-dependent calcium channels (Elliott and Ram, 2011). VER has been used in the treatment of cardiac vascular diseases such as cardiac arrhythmias, hypertension, and angina pectoris (Lido et al., 2022). VER has been reported to inhibit ROS production after ischemia–reperfusion of the rat liver (Jangholi et al., 2020) and demonstrated its efficacy in the treatment of oxidative stress-associated damage as a potent NRF2 activator (Lee et al., 2017).

We previously reported that only the combined (CURVER) and no the single treatment with CUR and VER significantly improved the cognitive phenotype of CD mice (Ortiz-Romero et al., 2021). Subsequently, we next aim was to evaluate if the combinatorial treatment could represent an effective pharmacological strategy to treat cardiovascular alterations in CD mice, as well as, the role of the XOR as another important source of oxidative stress in WBS. We here report that chronic combinatorial CURVER treatment, ameliorates cardiovascular injuries concomitantly with the normalization of XOR and 3-NT levels and increased levels of nuclear pNRF2.

## Results

### Only the combinatorial treatment with curcumin and verapamil prevents hypertension in CD animals

Systolic blood pressure (SYS) and heart beats per minute (BPM) were determined after four weeks of CUR, VER and combined CURVER treatments in conscious males and females mice using a Non-Invasive Blood Pressure System (Segura-Puimedon et al., 2014). Males and females were finally analyzed together once no significant sex differences were observed (Figure S1 A,B; Table S1).

SYS analysis showed a significant genotype/treatment interaction (F_3,131_=4.032, *p*=0.0088). In individual comparisons, we observed significant differences with respect to genotype in all CUR and VER experimental groups except for mice treated with CURVER, who presented differences with respect to treatment (*p*<0.0001) (Figure 1A).

**Figure 1:**
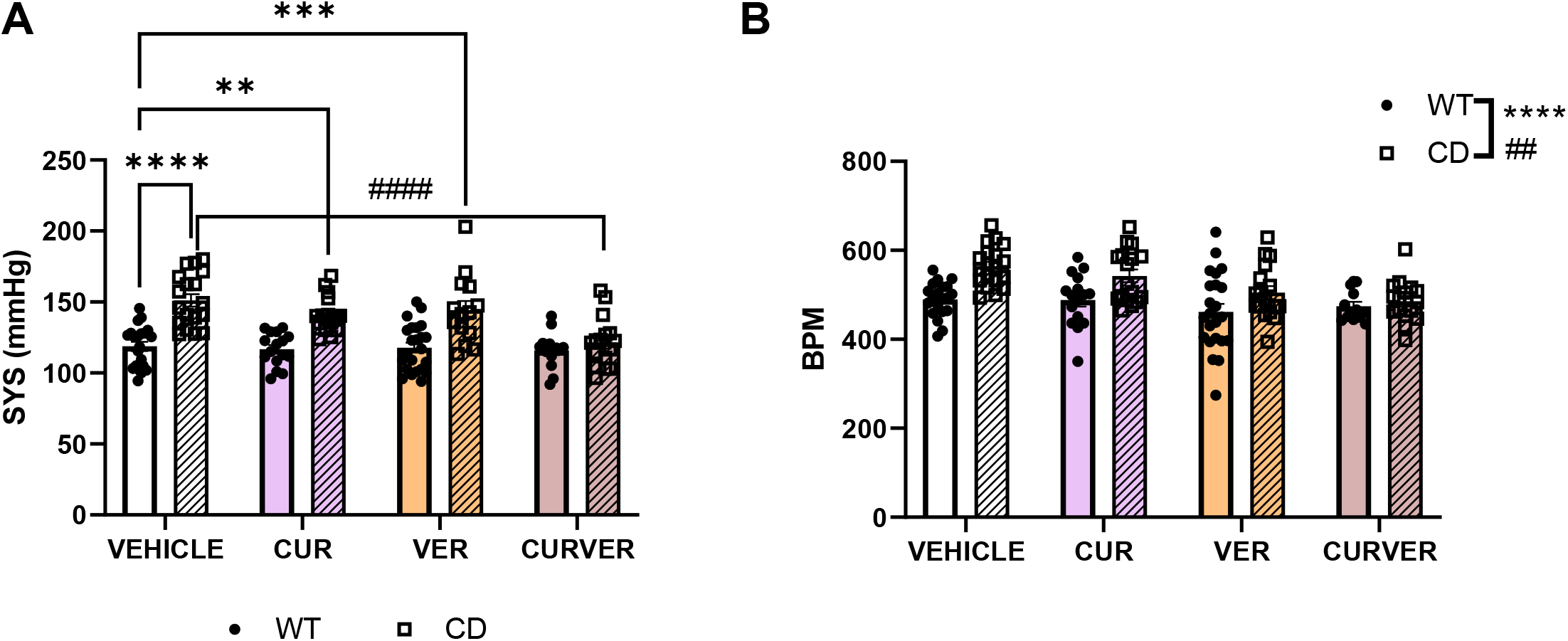
CURVER normalized systolic blood pressure and heart rate. **(A)** SYS blood pressure and **(B)** BPM measurements in the conscious state using the indirect tail-cuff method in WT and CD mice treated with single or combined treatments. A significant genotype/treatment interaction (F_3,131_=4.032, *p*=0.0088) was observed in SYS measurements. Data are represented as mean ± SEM. Two-way ANOVA with Tukey’s multiple comparisons test. * effect of genotype; #, effect of treatment. **,## p<0.005; ***,### p<0.001; ****,#### p< 0.0001

Interestingly, a two-way ANOVA analysis showed significant differences at the genotype level, with an increase in the number of BPM in CD animals compared to WT (F_1,133_=21.49, *p*<0.0001), in agreement with previous data that positively correlates the slopes of heart rate with basal blood pressure in the overall population (Marraccini et al., 1989). We also observed a significant effect of treatment (F_3,133_=5.677, *p*=0.0011) with a reduction in the heart rate of treated mice (Figure 1B).

None of treatments affected the WT either in SYS nor in BPM measures (Supplemental Figure S1 A,B, WT panels). Detailed statistical analysis is shown in Supplementary Table S1.

### Combinatorial treatment with curcumin and verapamil (CURVER) prevents aortic stenosis in CD mice

Considering above results, we next evaluated only CURVER administration experiments. Thus, with the aim of evaluating if the improvement in blood pressure in CURVER-treated CD mice correlates with improve of aortic parameters, we performed a histological comparison between the different groups.

In the ascending aortic wall of CD animals we observed a significant increase in the thickness of tunica media (effect of genotype: F_1, 30_ = 27.99, *p*<0.0001) (Figure 2A) together with a significant decrease of the aortic lumen diameter (effect of genotype F_1, 30_ = 42.17, *p*<0.0001) (Figure 2B). After CURVER treatment we observed a general improvement in lumen ascending aortic diaimeter and a reduction of the wall abnormal thickness (Figure 2A, B).

**Figure 2:**
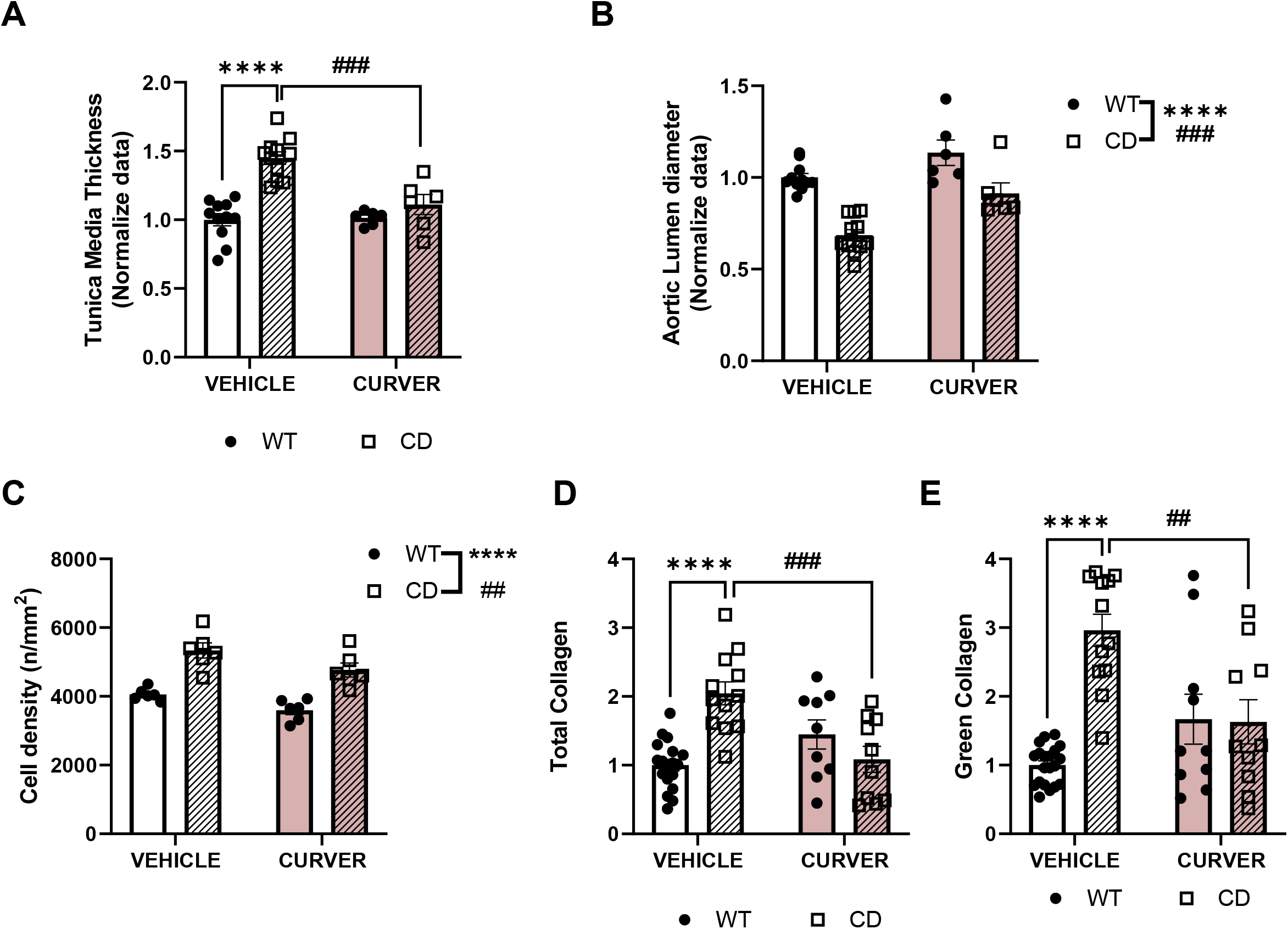
CURVER normalized stenosis in ascending aorta of treated CD mice. **(A)** Tunica media area thickness and **(B)** lumen diameter histograms comparisons. A significant genotype/treatment interaction (F_1,30_=11.60, *p*=0.0019) was observed in tunica media thickness. **(C)** Tunica media cell number by mm^2^. **(D)** Total collagen and **(E)** green collagen levels. All data has been normalized to WT values. Data are represented as mean ± SEM. Two-way ANOVA with Tukey’s multiple comparisons test. * effect of genotype; #, effect of treatment. ***,### p<0.001; ****,#### p< 0.0001

As expected, the amount of elastin in the tunica media of CD animals was significantly lower compared to WT animals (effect of genotype F_1, 30_ = 43.72, *p*<0.0001) regardless of CURVER treatment (F_1, 30_ = 0.5833, *p*=0.4510) (Figure S2A, B). We reasoned that the thickening of the tunica media observed in CD animals could be as a result of compensatory reaction increasing the number of VSMCs, of collagen content, or both. Thus, we next analyzed the density of SVMCs. We observed a significant increase of cell number in ascending aortas of CD mice (effect of genotype F_1,20_ = 53.93, *p*<0.0001). CURVER treatment significantly reduced the number of VSMCs (effect of treatment, F_1,20_ = 9.485, *p*= 0.0059) (Figure 2C).

Next, we analyzed the amount of collagen present in the tunica media by PicroSirius Red Staining. Under polarized light microscopy, green and red collagen fibers were observed, which are demonstrative of their different fiber thickness and assembly compaction, and, therefore, of their degree of maturity (Rittié, 2017). We observed a significant increase in total collagen fibers in the tunica media of CD mice (*p*<0.0001) (Figure 2D, Figure S2C), mainly due to a significant increase in immature (green) fibers (*p*<0.0001) (Figure 2E, Figure S2C). This remodeling was prevented after CURVER treatment with significant differences between vehicle and CURVER-treated CD mice (*p*=0.0005 for total and *p*=0.0015 for green) (Figure 2D,E and Figure S2C). In the case of mature (red) collagen fiber formation, a significant interaction between genotype/treatment was noted (F_1, 46_ = 8.411, *p*= 0.0057), although without reaching significant differences among groups (Figure S2D).

Detailed statistical analysis is shown in Supplementary Table S2.

### CURVER treatment attenuates the cardiopathy in CD mice

Cardiac hypertrophy was evaluated as the contribution of heart weight to total body weight. Males and females were analyzed together after verifying that there were no sex differences (Figure S3 and Table S3). The observed cardiac hypertrophy in CD mice (effect of genotype F_(1, 101)_ = 16.37, *p*=0.0001) was prevented by chronic CURVER administration, avoiding the increased heart weight (genotype/treatment interaction: F_(1,101)_=20.50, p<0.0001; Figure 3A). Histological analysis (Figure 3B) confirmed that CURVER treatment reduced (p=0.5491) the thickening of the left ventricular (LV) wall identified in vehicle-treated CD mice (p=0.0009) (Figure 2C).

**Figure 3:**
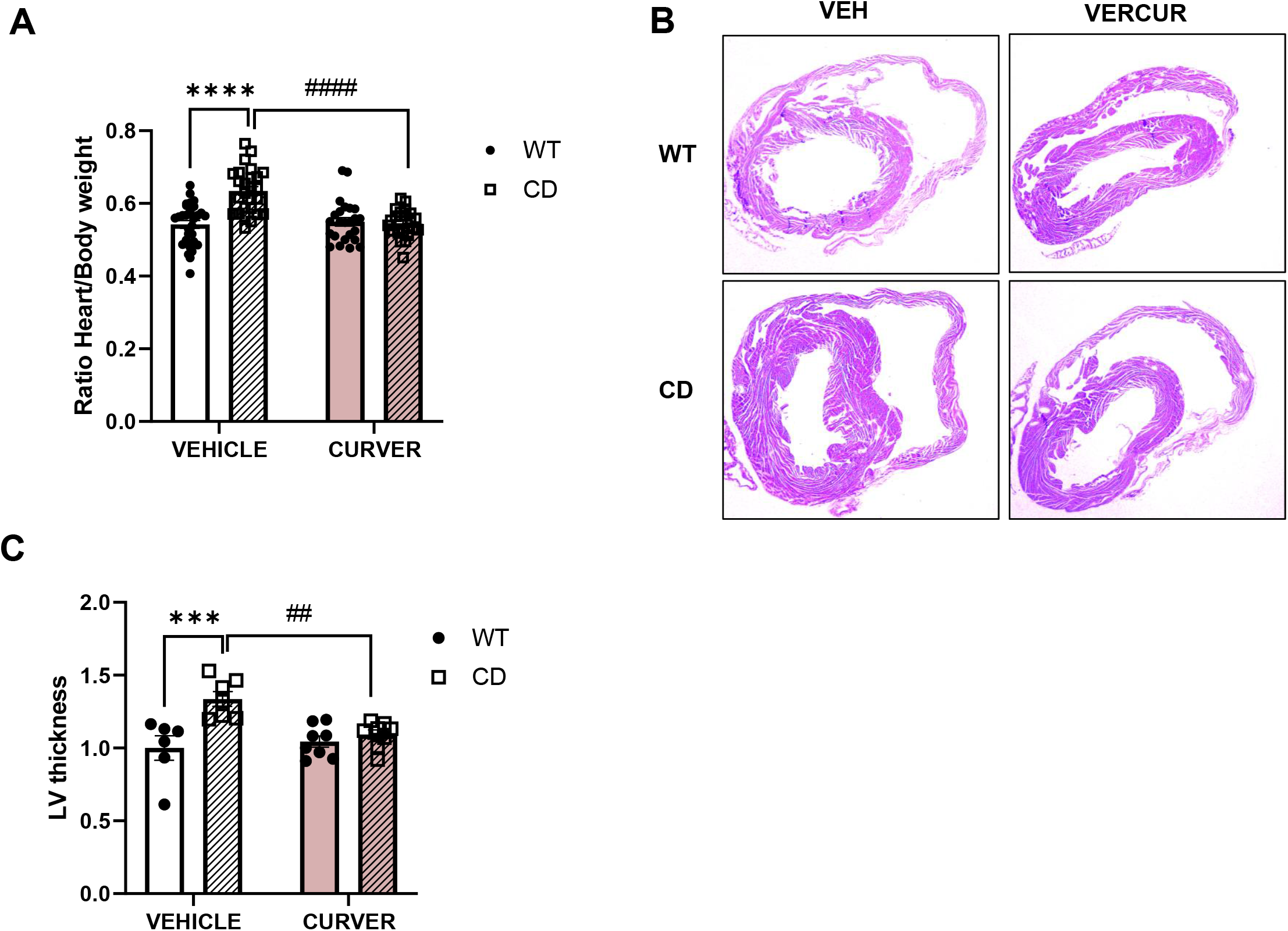
CURVER attenuates hyperthrophy in hearts of treated CD mice. **(A)** Significant cardiac hypertrophy (*p*<0.0001) was eliminated by chronic CURVER administration, preventing the enhanced heart weight (*p*=0.9995) **(B)** Representative images of histological analysis. **(C)** The significant thickening of the left ventricular wall identified in vehicle-treated CD mice (p=0.0009), was normalized with the CURVER treatment (p=0.5491). Data are represented as mean ± SEM. Two-way ANOVA with Tukey’s multiple comparisons test. * effect of genotype; #, effect of treatment. **,## p<0.005; ***,### p<0.001; ****,#### p< 0.0001

Detailed statistical analysis is shown in Supplementary Table S3.

### Concomitant decrease of XOR protein and 3’-nitrotyrosine residues levels after CURVER treatment

Focusing in oxidative stress, our main objective was to investigate the mechanism(s) by which CURVER treatment prevents the formation of the stenosis and cardiac hypertrophy. XOR, as an important source of ROS, has been found to play important roles in a variety of pathophysiological states in the cardiovascular system, including ischemia/reperfusion injury, atherosclerosis, and left ventricular dysfunction after myocardial infarction (Kinugasa et al., 2003; Engberding et al., 2004). Endothelial cell-generated nitric oxide (NO) easily interacts with XOR-derived anion superoxide (O^-^_2_) forming peroxynitrite (ONOO^-^), which in the endothelium of the tunica intima and VSMCs of the media irreversibly generates reactive nitrogen species (RNS) residues such as 3’-nitrotyrosine (3-NT) (Houston et al., 1998; Takata et al., 2014), which is usually used as a redox stress marker.

Thus, we evaluated first the levels of the redox stress by quantifying the levels of protein nitrosylation in the ascending aortas wall. CD mice showed significantly higher levels of 3 NT when compared to WT littermates (*p*=0.0008) (Figure 4A). These increased levels could be associated to the concomitant observed increase of protein XOR levels (*p*=0.0478). Our data demonstrated that after the combined CURVER treatment, CD mice showed significant decrease of 3-NT and XOR aortic levels (interaction genotype/treatment F_(1, 28)_ = 11.27, *p*=0.0023 and F_(1, 19)_ = 5.952, *p*=0.0247, respectively) (Figure 4 A,B).

**Figure 4:**
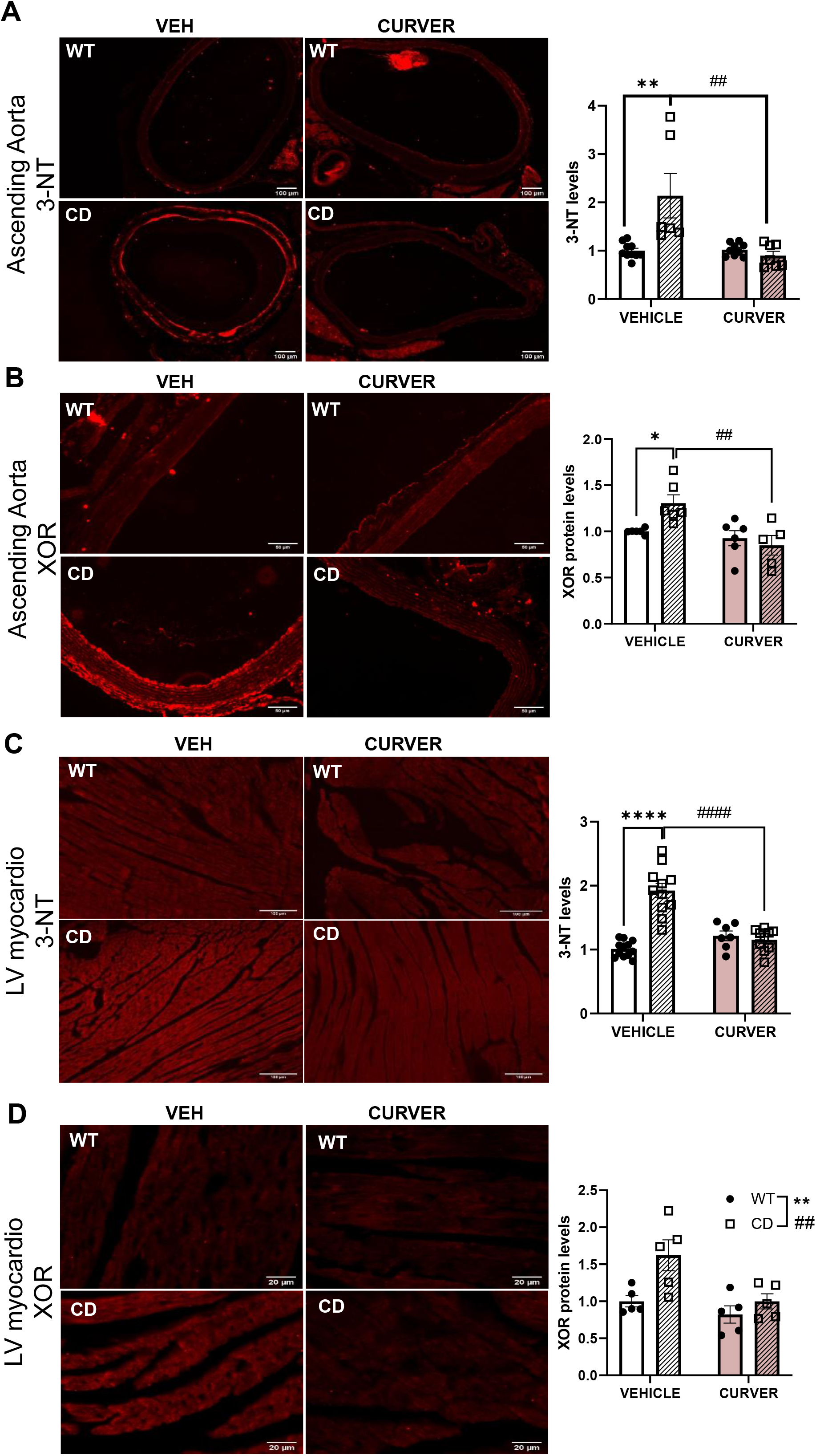
Concomitant reduction of 3-NT and XOR levels after CURVER treatment. Representative images of 3-NT immunodetection in ascending aortas **(A)**, quantification showed a significant interaction genotype/treatment F_(1, 28)_ = 12.12, *p*=0.0017). **(B)** Significant increase in the levels of XOR could be appreciated in CD vehicle ascending aorta that were decreased in CURVER treated CD mice (interaction genotype/treatment F_(1, 19)_ = 5.952, *p*=0.0247) In heart, we observed a significant interaction genotype/treatment F_(1, 36)_ = 39,09 *P<*0.0001) in the 3-NT levels **(C)** together with a significant increase in the levels of XOR of CD mice (effect of genotype, F_(1, 16)_ =8.653, *p*=0.0096) that was reduced after CURVER treatment (effect of treatment, F_(1, 16)_ = 8.725, *p*=0.0093) **(D)**. Data are represented as mean ± SEM. Two-way ANOVA with Tukey’s multiple comparisons test. * effect of genotype; #, effect of treatment. *,# p<0.05; **,## p<0.005; ***,### p<0.001; ****,#### p< 0.0001

Similar results were observed in LV-myocardium, with increased levels of 3-NT (effect of genotype, F_(1, 36)_ = 29.96, *p*<0.0001) (Figure 4C) and XOR (effect of genotype, F_(1, 16)_ = 8.653, *p*=0.0096) (Figure 4D) that were reduced after CURVER treatment (effect of treatment F_(1, 36)_ = 13.10, *p*=0.0009 in 3-NT and F_(1, 16)_ = 8.725, *p*=0.0093 for XOR) (Figure 4 C,D). XOR protein levels were validated by Western blot (Figure S4A, B). Detailed statistical data are shown in Supplemental Table S4.

### CURVER treatment stimulates the nuclear translocation of pNRF2 in CD mice

Previous studies have shown that nuclear levels of pNRF2 were decreased in cultured primary CD cardiomyocytes with the corresponding decreased expression levels of targets endogenous antioxidants genes as *Nqo1* (Ortiz-Romero et al., 2018). Both CUR and VER have been described as potent NRF2 activators (Zeng et al., 2015; Lee et al., 2017). In this way, we analyzed the effect of treatment in the pNRF2 levels in ascending aortas and L-myocardium.

Inmunofluorescence analysis of the tunica media of ascending aortas showed a significant reduced number of pNFR2 positive cells in vehicle treated-CD mice (P<0.0001) that normalized after CURVER treatment (P=0.5890) (Figure 5A). In agreement with previous results, we observed a significant reduction in pNRF2 levels in total protein LV-myocardium extract of CD mice compared to WT (P=0.0084). After CURVER treatment CD-treated mice showed a significant increase of pNRF2 respect to CD-vehicle mice (P= 0.002)(Figure 5B).

**Figure 5:**
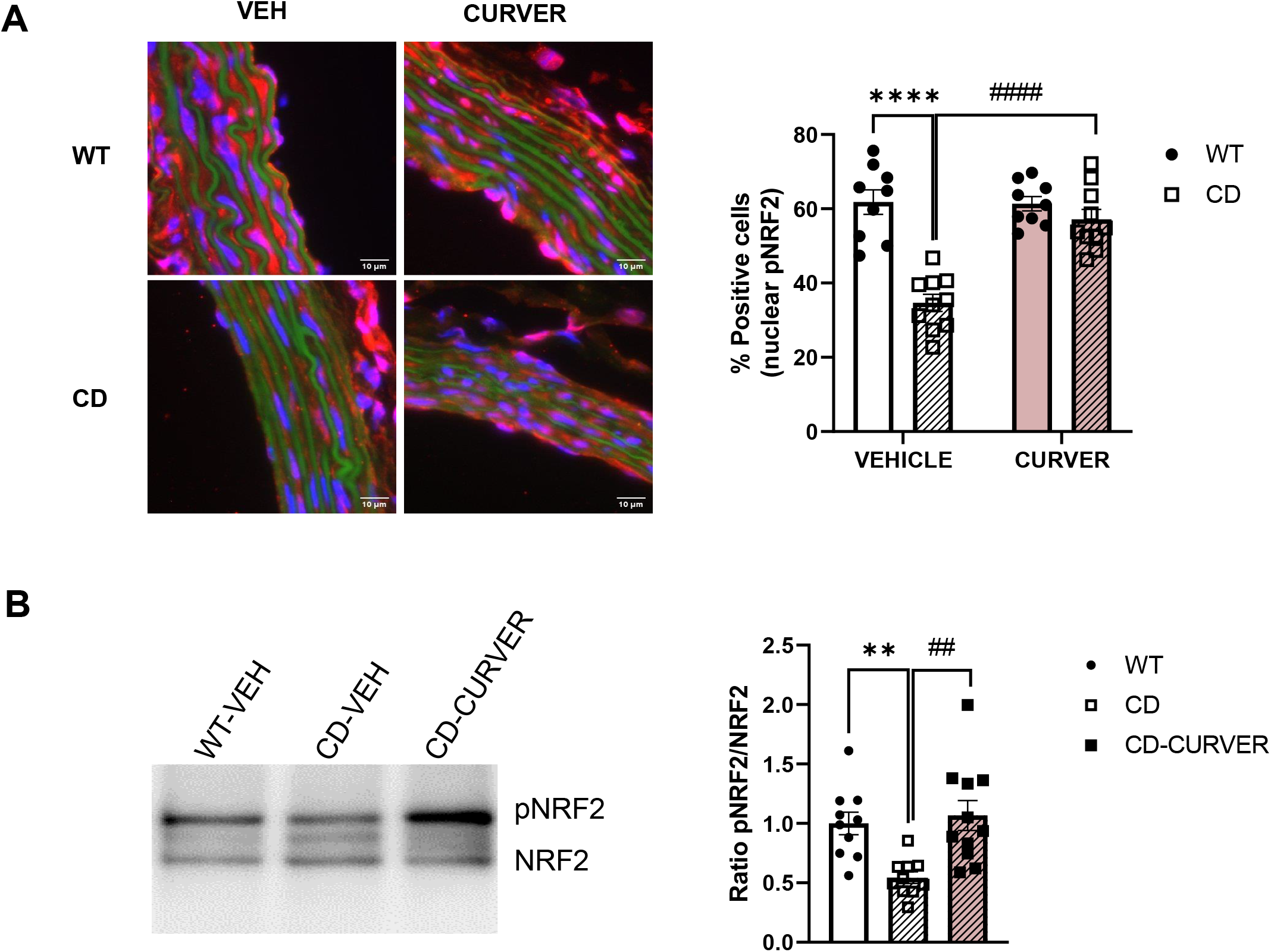
CURVER increased pNRF2 nuclear location in the cardiovascular system of CD mice. (A) Inmunohistofluorescence was performed to detect the pNRF2 in SVMCs of the tunica media (red) nuclear staining (DAPI) in blue (Left panel). Significant genotype/treatment interaction could be observed (F_(1, 34)_ = 19,30; *P*=0,0001). Two-way ANOVA with Tukey’s multiple comparisons test **(B)** Western blot (representative image, left) shown significant reduction in the pNRF2 levels in LV-myocardium protein extract of CD mice compared to WTs (P=0.0084). After CURVER treatment CD-treated mice showed a significant increase of pNRF2 respect to CD-vehicle mice (P= 0.002). One way ANOVA, followed by Tukey’s post hoc comparisons. Data are represented as mean ± SEM.. * effect of genotype; #, effect of treatment. **,## p<0.005; ***,### p<0.001; ****,#### p< 0.0001

Detailed statistical analysis is shown in Supplementary Table S5.

## Discussion

In this work we analyzed the effect of the combined treatment curcumin, the most abundant phenol in turmeric, and verapamil, a widely used medication, on the cardiovascular phenotype of CD murine model of WBS.

Hypertension is a consistent finding among WBS patients. In a comparison of WBS individuals with control population hypertension was present in 40% of adult patients versus 14% of controls. The difference was even more dramatic in infantile patients, with 46% of them presenting hypertension versus 6% of controls (Broder et al., 1999). Curcumin and verapamil are able by themselves of improving arterial hypertension (Schmieder et al., 1987; Yao et al., 2016). In this study, both compounds had this effect and the individual treatments caused a reduction of systolic pressure by 10% in CD animals. However, this reduction was not enough to reach WT levels of systolic pressure, and only the combinatorial treatment with both compounds was able to achieve this significant changes. In relation to the heart rate, we observed an effect of the treatment, although without interaction with the genotype. A detailed analysis of data shows an absence of effect of curcumin, while we observed a small decrease in heart rate in animals treated with verapamil, regardless of genotype. Unlike in WT mice, the combined treatment increases the reduction observed with verapamil alone in CD mice.

The reduction in hypertension was accompanied by an improvement in the histological parameters studied, both at the level of ascending aortic stenosis and cardiac hypertrophy. We also report for the first time, a significant increase in immature collagen fibers present in the tunica media of the ascending aortas of CD animals. In accordance with previous studies implicating verapamil with collagen synthesis (Zeng et al., 2021), and the role of curcumin in better maturation and cross linking of collagen (Panchatcharam et al., 2006), after CURVER treatment the increase in immature fibers observed in untreated CD mice was significantly reduced. This process could contribute to the remodeling of the ascending aorta, reducing the thickness of the tunica media and augmenting the lumen diameter. In parallel with the improvement in aortic stenosis, we observed a reduction in cardiac hypertrophy, with a normalization of the thickness of LV-myocardium wall.

Other main objective of our study was to obtain more information about the contribution of redox stress in the molecular pathology of the cardiovascular phenotype of WBS paying special attention to XOR, which, like NADPH oxidases, is another important source of ROS in the cardiovascular system (Polito et al., 2021). In this study we have shown high levels of XOR protein both in the ascending aorta and in cardiac tissue. These high levels correlate with increased levels of 3-NT, which are especially visible in the endothelial cells (tunica intima). This increase in 3-NT in the aortic intima of WBS is consistent with the recent demonstration of increased nNOS in aortic stenosis formation in CD animals (Jiménez-Altayó et al., 2020). In CURVER treated CD mice we observed a significant reduction of XOR protein levels both in ascending aorta and LV myocardium, in agreement in previous data (Chen et al., 2019). The reduction of XOR protein levels correlates with reduction of 3-NT staining.

In this line of evidence, we previously reported that redox stress present in cardiac tissue seems not to be attenuated due to the intrinsic physiological antioxidant response induced by the nuclear translocation of pNRF2 (Ortiz-Romero et al., 2018). We demonstrated that this response was also altered in the tunica media of ascending aorta, with a significant reduction of nuclear pNRF2 positive cells in CD mice. An increase in the nuclear localization of pNRF2, with the concomitant increase in the expression of some antioxidant ARE response genes after treatment could be observed in CD-treated mice. Of note, the induction of NRF2 had already been described previously in the case of CUR and VER (Zeng et al., 2015; Lee et al., 2017).

In conclusion, a combination of curcumin and verapamil significantly ameliorates the cardiovascular phenotype of a mouse model for WBS. Its efficiency is achieved through reduction of oxidative stress levels evidenced by the reduction of XOR protein levels and induction of NRF2 pathway, both in the aortic wall and LV-myocardium. Verapamil is already approved for human use, while curcumin is a natural safe product. We here shown that their combination deserves further evaluation as potential therapeutic agent to prevent the severe cardiovascular injuries occurring in human patients with WBS.

## MATERIALS AND METHODS

### Ethics statement

The local Committee of Ethical Animal Experimentation (CEEA-PRBB; Protocol Number: VCU-17-0021; and University of Barcelona Protocol Number: Campuzano-153.19) and Generalitat de Catalunya (Protocol Number: DAAM-9494) approved all animal procedures in accordance with the guidelines of the European Communities Directive 86/609/EEC. The PRBB has Animal Welfare Assurance (#A5388-01, Institutional Animal Care and Use Committee approval date 05/08/2009), granted by the Office of Laboratory Animal Welfare (OLAW) of the US National Institutes of Health.

### Animals’ maintenance

CD mice, a WBS murine model that carries a 1.3 Mb heterozygous deletion harboring from *Gtf2i* to *Fkbp6*, were obtained as previously described (Segura-Puimedon et al., 2014). All mice were maintained on 97% C57BL/6J background. Genomic DNA was extracted from mouse ear punch to perform the genotyping using PCR and appropriate primers, as previously described (Borralleras et al., 2015). Animals were housed under standard conditions in a 12 h dark/light cycle with access to food and water/treatment *ad libitum*.

### Treatment administration and intake

The treatment was administered following the protocols previously used by the group (Ortiz-Romero et al., 2021). Vehicle control group (VEH) consisted of WT and CD mice drinking water with 1% DMSO. Experimental groups consisted of: 1) WT and CD mice drinking verapamil (VER); 2) WT and CD mice drinking curcumin (CUR) and 3) WT and CD mice drinking the combination of both compounds (CURVER). Treatment was started at 8 weeks-old (young mice). All animals drank their treatments for 1 month before measuring arterial pressure, and then sacrificed at 12 weeks-old (young adult).

Two-way ANOVA of repeated measures shown not significant interaction between time and treatment (Males: F_45,240_ = 0,7963, *p*=0.8189; Females: F_45, 240_ = 0,7335, *p*=0.8936) (Figure S6A). There were also no significant differences in the consumption of the treatments in relation to the genotype (F_3, 24_ = 0.1347, *p*=0.9384) (Figure S6B).

In agreement with previous reports (Segura-Puimedon et al., 2014) CD mice, both males and females, had significantly lower body weights compared to WT mice (effect of genotype F_1,102_ = 92.76, p<0.0001 in males; F_1,62_ = 38.57, p<0.0001 in females), without significant effect of treatment (Figure S6 C, D). Supplemental Table S6 shows detailed statistical analysis.

### Heart rate and Arterial pressure measurement

Beats by minute (BPM) and Systolic blood pressure (SYS) of conscious mice were measured using a tail cuff system (Non-Invasive Blood Pressure System, PanLab), while holding mice in a restrainer tube that was cleaned after each mouse. After habituation to the restrainer, measures were taken on three separate occasions and averaged for each mouse.

### Histological preparation

Animals were perfused with 1X phosphate buffered saline (PBS) followed by 4% paraformaldehyde (PFA). Hearts and aortas were removed and post-fixed in 4% PFA for 48 h at 4 °C and conserved in 75% EtOH until the generation of the paraffin blocks. Paraffin-embedded hearts and tissue arrays of mice aortae (TMAs) from different experimental sets were cut in 5 μm sections.

All measurements were carried out in a blinded manner by two different observers with no knowledge of genotype and treatment. Images were captured using a Leica Leitz DMRB microscope (10x objective) equipped with a Leica DC500 camera and analyzed with Fiji Image J Analysis software. For collagen analysis were performed with PicroSirius Red Staining method and analyzed with a predesigned macro program (Rubies et al., 2019).

### Immunofluorescence

Paraffin-embedded cardiac and aortic tissue sections were deparaffinized and rehydrated prior to unmasking the epitope. To unmask epitopes, tissue sections were treated with a retrieval solution (10 mM sodium citrate, 0.05% Tween, pH 6) for 30 min in the steamer at 95°C. Next, sections were incubated for 20 minutes with ammonium chloride (NH4Cl, 50 mM, pH 7.4) to block free aldehyde groups, followed by permeabilization step (0,3% Triton x-100; 10 min), rinsed three times with PBS and then incubated with 1% BSA in PBS for 2 hours at RT prior to overnight incubation in humidified chamber at 4°C with the corresponding primary antibody; anti-3NT (1:200; Merck Millipore 06-284), anti-XOR (1:50; Rockland 200-4183S) and, anti-pNRF2 (1:200; Abcam ab76026).On the next day, sections were incubated with secondary antibodies solution anti-rabbit, Alexa 555, Alexa 647 (1:2.000, Invitrogen). Finally, sections were stained with DAPI and mounted with fluorescence mounting medium (CAT NO 0100-01; Southern Biotech) over the same glass slide.

For quantitative analysis of immunostainings, four areas of each corresponding tissue section were quantified with Image J software. All measurements were carried out in a blinded manner by two independent investigators.

### Western blotting

Dissected tissues were homogenized with bullet blender and beads in RIPA buffer containing protease inhibitors (2 mM phenylmethylsulphonyl fluoride, 10 g/L aprotinin, and 1 g/L leupeptin, 1 g/L pepstatin), and phosphatase inhibitors (2 mM Na3VO4 and 100 mM NaF). Protein concentration was determined using the Dc protein assay kit (Bio-Rad). Membranes were blotted overnight at 4°C with the following primaries antibodies: anti-XDH (1:500 Santa Cruz sc-398548), anti-NRF2 (1:500 Santa Cruz sc-H300) and pNRF2 (1:5000, Abcam ab). Later, membranes were washed and incubated with the appropriate HRP conjugated secondary antibody (1:3,000 anti-rabbit W401B antibody from Promega), and the reaction was finally visualized with the Western Blotting Luminol Reagent (Santa Cruz Biotechnology). Western blot replicates were quantified using a computer-assisted densitometric analysis (Gel-Pro Analyzer, version 4; Media Cybernetics).

### Statistical analysis

Prior to analysis all data were analyzed by the Shapiro-Wilk test to confirm normality. Males and females were analyzed together after verifying that there were no gender differences for the analyzed parameter (three way ANOVA, gender-genotype-treatment). One-two- or three way ANOVA with Tukey or Šídák’s *post hoc* test were used when needed. All data are presented as mean ± SEM. Values were considered significant when *p*<0.05. GraphPad Prism 9 software was used for obtaining all statistical tests and graphs.

## Acknowledgements

We thank Maria Encarnación Palomo, and the Advanced Microscopy Unit of the University of Barcelona for Technical assistance. We thank Montserrat Batlle for helping in the Sirius red staining and analysis.

## Author Contributions

Substantial contributions to the conception/design of the work: GE-LPJ-VC

The acquisition, analysis, drafting and interpretation of data for the work: NA-POR-IRR-VC

Revising the work critically for important intellectual content: GE-LPJ-VC

Final approval of the version to be published: NA-POR-IRR-GE-LPJ-VC

Agreement to be accountable for all aspects of the work in ensuring that questions related to the accuracy or integrity of any part of the work are appropriately investigated and resolved: NA-POR-IRR-GE-LPJ-VC.

## References

Akter, J., Islam, Z., Hossain, A., and Takara, K. (2020). Pharmacological activities of 4-methylene-8-hydroxybisabola-2,10-diene-9-one, a new compound isolated from Ryudai gold (Curcuma longa). Naunyn. Schmiedebergs. Arch. Pharmacol. 393, 191–201. doi:10.1007/s00210-019-01721-3.

Borralleras, C., Sahun, I., Pérez-Jurado, L. A., and Campuzano, V. (2015). Intracisternal Gtf2i gene therapy ameliorates deficits in cognition and synaptic plasticity of a mouse model of williams-beuren syndrome. Mol. Ther. 23. doi:10.1038/mt.2015.130.

Bozkurt, O., Kocaadam-Bozkurt, B., and Yildiran, H. (2022). Effects of curcumin, a bioactive component of turmeric, on type 2 diabetes mellitus and its complications: an updated review. Food Funct. doi:10.1039/d2fo02625b.

Broder, K., Reinhardt, E., Ahern, J., Lifton, R., Tamborlane, W., and Pober, B. (1999). Elevated ambulatory blood pressure in 20 subjects with Williams syndrome. Am. J. Med. Genet. 83, 356–60. doi:10.1002/(sici)1096-8628(19990423)83:5<356::aid-ajmg2>3.0.co;2-x.

Campuzano, V., Segura-Puimedon, M., Terrado, V., Sánchez-Rodríguez, C., Coustets, M., Menacho-Márquez, M., et al. (2012). Reduction of NADPH-oxidase activity ameliorates the cardiovascular phenotype in a mouse model of Williams-Beuren Syndrome. PLoS Genet. 8. doi:10.1371/journal.pgen.1002458.

Chen, Y., Li, C., Duan, S., Yuan, X., Liang, J., and Hou, S. (2019). Curcumin attenuates potassium oxonate-induced hyperuricemia and kidney inflammation in mice. Biomed. Pharmacother.118, 109195. doi:10.1016/j.biopha.2019.109195.

Costa, T. J., Barros, P. R., Arce, C., Santos, J. D., da Silva-Neto, J., Egea, G., et al. (2021). The homeostatic role of hydrogen peroxide, superoxide anion and nitric oxide in the vasculature. Free Radic. Biol. Med. 162, 615–635. doi:10.1016/j.freeradbiomed.2020.11.021.

Del Campo, M., Antonell, A., Magano, L. F., Muñoz, F. J., Flores, R., Bayés, M., et al. (2006). Hemizygosity at the NCF1 gene in patients with Williams-Beuren syndrome decreases their risk of hypertension. Am. J. Hum. Genet. 78, 533–542. doi:10.1086/501073.

du Preez, R., Pahl, J., Arora, M., Ravi Kumar, M. N. V., Brown, L., Panchal, S. K., et al. (2019). Low-Dose Curcumin Nanoparticles Normalise Blood Pressure in Male Wistar Rats with Diet-Induced Metabolic Syndrome. Nutrients 11, 1542. doi:10.3390/nu11071542.

Elliott, W. J., and Ram, C. V. S. (2011). Calcium Channel Blockers. J. Clin. Hypertens. 13, 687–689. doi:10.1111/j.1751-7176.2011.00513.x.

Engberding, N., Spiekermann, S., Schaefer, A., Heineke, A., Wiencke, A., Müller, M., et al. (2004). Allopurinol attenuates left ventricular remodeling and dysfunction after experimental myocardial infarction: a new action for an old drug? Circulation 110, 2175–9. doi:10.1161/01.CIR.0000144303.24894.1C.

Houston, M., Chumley, P., Radi, R., Rubbo, H., and Freeman, B. A. (1998). Xanthine Oxidase Reaction with Nitric Oxide and Peroxynitrite. Arch. Biochem. Biophys. 355, 1–8. doi:10.1006/abbi.1998.0675.

Jangholi, E., Sharifi, Z. N., Hoseinian, M., Zarrindast, M.-R., Rahimi, H. R., Mowla, A., et al. (2020). Verapamil Inhibits Mitochondria-Induced Reactive Oxygen Species and Dependent Apoptosis Pathways in Cerebral Transient Global Ischemia/Reperfusion. Oxid. Med. Cell. Longev. 2020, 1–12. doi:10.1155/2020/5872645.

Jiménez-Altayó, F., Ortiz-Romero, P., Puertas-Umbert, L., Dantas, A. P. A. P., Pérez, B., Vila, E., et al. (2020). Stenosis coexists with compromised α1-adrenergic contractions in the ascending aorta of a mouse model of Williams-Beuren syndrome. Sci. Rep. 10, 1–12. doi:10.1038/s41598-020-57803-3.

Kinugasa, Y., Ogino, K., Furuse, Y., Shiomi, T., Tsutsui, H., Yamamoto, T., et al. (2003). Allopurinol improves cardiac dysfunction after ischemia-reperfusion via reduction of oxidative stress in isolated perfused rat hearts. Circ. J. 67, 781–7. doi:10.1253/circj.67.781.

Kozel, B. A., Danback, J. R., Waxler, J. L., Knutsen, R. H., de Las Fuentes, L., Reusz, G. S., et al. (2014). Williams syndrome predisposes to vascular stiffness modified by antihypertensive use and copy number changes in NCF1. Hypertension 63, 74–79. doi:10.1161/HYPERTENSIONAHA.113.02087.

Lee, D. H., Park, J. S., Lee, Y. S. Y. ho, Sung, S. H., Lee, Y. S. Y. ho, and Bae, S. H. (2017). The hypertension drug, verapamil, activates Nrf2 by promoting p62-dependent autophagic Keap1 degradation and prevents acetaminophen-induced cytotoxicity. BMB Rep. 50, 91–96. doi:10.5483/BMBRep.2017.50.2.188.

Liang, J., Yamaguchi, Y., Matsumura, F., Goto, M., Akizuki, E., Matsuda, T., et al. (2000). Calcium-channel blocker attenuates Kupffer cell production of cytokine-induced neutrophil chemoattractant following ischemia-reperfusion in rat liver. Dig. Dis. Sci. 45, 201–9. doi:10.1023/a:1005498402659.

Lido, P., Romanello, D., Tesauro, M., Bei, A., Perrone, M. A., Palazzetti, D., et al. (2022). Verapamil: prevention and treatment of cardio-renal syndromes in diabetic hypertensive patients? Eur. Rev. Med. Pharmacol. Sci. 26, 1524–1534. doi:10.26355/eurrev_202203_28217.

Llano, S., Gómez, S., Londoño, J., and Restrepo, A. (2019). Antioxidant activity of curcuminoids. Phys. Chem. Chem. Phys. 21, 3752–3760. doi:10.1039/c8cp06708b.

Marraccini, P., Palombo, C., Giaconi, S., Michelassi, C., Genovesi-Ebert, A., Marabotti, C., et al. (1989). Reduced cardiovascular efficiency and increased reactivity during exercise in borderline and established hypertension. Am. J. Hypertens. 2, 913–6. doi:10.1093/ajh/2.12.913.

Ortiz-Romero, P., Borralleras, C., Bosch-Morató, M., Guivernau, B., Albericio, G., Muñoz, F. J. F. J., et al. (2018). Epigallocatechin-3-gallate improves cardiac hypertrophy and short-term memory deficits in a Williams-Beuren syndrome mouse model. PLoS One 13, 1–19. doi:10.1371/journal.pone.0194476.

Ortiz-Romero, P., González-Simón, A., Egea, G., Pérez-Jurado, L. A., and Campuzano, V. (2021). Co-Treatment With Verapamil and Curcumin Attenuates the Behavioral Alterations Observed in Williams-Beuren Syndrome Mice by Regulation of MAPK Pathway and Microglia Overexpression. Front. Pharmacol. 12, 670785. doi:10.3389/fphar.2021.670785.

Panchatcharam, M., Miriyala, S., Gayathri, V. S., and Suguna, L. (2006). Curcumin improves wound healing by modulating collagen and decreasing reactive oxygen species. Mol. Cell. Biochem. 290, 87–96. doi:10.1007/s11010-006-9170-2.

Polito, L., Bortolotti, M., Battelli, M. G., and Bolognesi, A. (2021). Xanthine oxidoreductase: A leading actor in cardiovascular disease drama. Redox Biol. 48, 102195. doi:10.1016/j.redox.2021.102195.

Rittié, L. (2017). Method for Picrosirius Red-Polarization Detection of Collagen Fibers in Tissue Sections. Methods Mol. Biol. 1627, 395–407. doi:10.1007/978-1-4939-7113-8_26.

Rodríguez-Rovira, I., Arce, C., De Rycke, K., Pérez, B., Carretero, A., Arbonés, M., et al. (2022). Allopurinol blocks aortic aneurysm in a mouse model of Marfan syndrome via reducing aortic oxidative stress. Free Radic. Biol. Med. 193, 538–550. doi:10.1016/j.freeradbiomed.2022.11.001.

Rubies, C., Dantas, A.-P., Batlle, M., Torres, M., Farre, R., Sangüesa, G., et al. (2019). Aortic remodelling induced by obstructive apneas is normalized with mesenchymal stem cells infusion. Sci. Rep. 9, 11443. doi:10.1038/s41598-019-47813-1.

Schmieder, R. E., Messerli, F. H., Garavaglia, G. E., and Nunez, B. D. (1987). Cardiovascular effects of verapamil in patients with essential hypertension. Circulation 75, 1030–6. doi:10.1161/01.cir.75.5.1030.

Segura-Puimedon, M., Sahún, I., Velot, E., Dubus, P., Borralleras, C., Rodrigues, A. J. A. J., et al. (2014). Heterozygous deletion of the Williams-Beuren syndrome critical interval in mice recapitulates most features of the human disorder. Hum. Mol. Genet. 23, 6481–6494. doi:10.1093/hmg/ddu368.

Senoner, T., and Dichtl, W. (2019). Oxidative Stress in Cardiovascular Diseases: Still a Therapeutic Target? Nutrients 11. doi:10.3390/nu11092090.

Soto, M. E., Manzano-Pech, L. G., Guarner-Lans, V., Díaz-Galindo, J. A., Vásquez, X., Castrejón-Tellez, V., et al. (2020). Oxidant/Antioxidant Profile in the Thoracic Aneurysm of Patients with the Loeys-Dietz Syndrome. Oxid. Med. Cell. Longev. 2020, 1–17. doi:10.1155/2020/5392454.

Takata, M., Amiya, E., Watanabe, M., Omori, K., Imai, Y., Fujita, D., et al. (2014). Impairment of flow-mediated dilation correlates with aortic dilation in patients with Marfan syndrome. Heart Vessels 29, 478–485. doi:10.1007/s00380-013-0393-3.

Yao, Y., Wang, W. E., Li, M., Ren, H., Chen, C., Wang, J., et al. (2016). Curcumin Exerts its Anti-hypertensive Effect by Down-regulating the AT1 Receptor in Vascular Smooth Muscle Cells. Sci. Rep. 6, 25579. doi:10.1038/srep25579.

Zeng, C., Zhong, P., Zhao, Y., Kanchana, K., Zhang, Y., Khan, Z. A., et al. (2015). Curcumin protects hearts from FFA-induced injury by activating Nrf2 and inactivating NF-κB both in vitro and in vivo. J. Mol. Cell. Cardiol. 79, 1–12. doi:10.1016/j.yjmcc.2014.10.002.

Zeng, M.-Q., Xiao, W., Yang, K., Gao, Z.-Y., Wang, J.-S., Lu, Q., et al. (2021). Verapamil inhibits ureteral scar formation by regulating CaMK II-mediated Smad pathway. Chem. Biol. Interact.346, 109570. doi:10.1016/j.cbi.2021.109570.

