## Supplemental Data for "Curcumin combined with verapamil improve cardiovascular phenotype of a Williams-Beuren Syndrome mice model reducing oxidative stress"

#### Supplemental Table S1: Statistical data of SYS and BPM

| Three-way ANOVA Sex effect SYS | F (DFn, DFd) | P value |
|---|---|---|
| Treatment | F (3, 123) = 4.692 | P=0.0039 |
| Genotype | F (1, 123) = 57.05 | P<0.0001 |
| Sex | F (1, 123) = 3.103 | P=0.0807 |
| Treatment x Genotype | F (3, 123) = 3.819 | P=0.0117 |
| Treatment x Sex | F (3, 123) = 0.6550 | P=0.5813 |
| Genotype x Sex | F (1, 123) = 0.9970 | P=0.3200 |
| Treatment x Genotype x Sex | F (3, 123) = 0.1893 | P=0.9035 |

| Three-way ANOVA sex effect BPM | F (DFn, DFd) | P value |
|---|---|---|
| Treatment | F (3, 125) = 5.543 | P=0.0013 |
| Genotype | F (1, 125) = 19.71 | P<0.0001 |
| Sex | F (1, 125) = 0.4535 | P=0.5019 |
| Treatment x Genotype | F (3, 125) = 1.623 | P=0.1873 |
| Treatment x Sex | F (3, 125) = 0,2480 | P=0.8627 |
| Genotype x Sex | F (1, 125) = 1.456 | P=0.2299 |
| Treatment x Genotype x Sex | F (3, 125) = 0.5289 | P=0.6632 |

| Two-way ANOVA SYS | F (DFn, DFd) | P value |
|---|---|---|
| Interaction | F (3, 131) = 4.032 | P=0.0088 |
| Treatment | F (3, 131) = 5.777 | P=0.0010 |
| Genotype | F (1, 131) = 62.25 | P<0.0001 |
| **Tukey's multiple comparisons test** | | **Adjusted P Value** |
| **VEHI:WT vs. VEH:CD** | | <0.0001 |
| VEH:WT vs. VER:WT | | >0.9999 |
| **VEH:WT vs. VER:CD** | | 0.0003 |
| VEH:WT vs. CUR:WT | | >0.9999 |
| **VEH:WT vs. CUR:CD** | | 0.0013 |
| VEH:WT vs. CURVER:WT | | >0.9997 |
| VEH:WT vs. CURVER:CD | | >0.9995 |
| VEH:CD vs. VER:WT | | <0.0001 |
| VEH:CD vs. VER:CD | | 0.9381 |
| VEH:CD vs. CUR:WT | | <0.0001 |
| VEH:CD vs. CUR:CD | | 0.6030 |

| | |
|---|---|
| VEH:CD vs. CURVER:WT | <0.0001 |
| **VEH:CD vs. CURVER:CD** | <0.0001 |

| **Two-way ANOVA BPM** | **F (DFn, DFd)** | **P value** |
|---|---|---|
| Interaction | F (3, 133) = 1,692 | P=0,1718 |
| Treatment | F (3, 133) = 5,677 | P=0,0011 |
| Genotype | F (1, 133) = 21,49 | P<0,0001 |

**Supplemental Table S2: Statistical data of Ascending Aorta parameters**

| **ANOVA thickness T. Media** | **F (DFn, DFd)** | **P value** |
|---|---|---|
| Interaction | F (1, 30) = 11,60 | P=0,0019 |
| Treatment | F (1, 30) = 9,898 | P=0,0037 |
| Genotype | F (1, 30) = 27,99 | P<0,0001 |
| **Tukey's multiple comparisons test** | | **Adjusted P Value** |
| **VEHICLE:WT vs. VEHICLE:CD** | | <0,0001 |
| VEHICLE:WT vs. CURVER:CD | | 0,4408 |
| **VEHICLE:CD vs. CURVER:CD** | | 0,0004 |

| **Two-way ANOVA diameter Lumen** | **F (DFn, DFd)** | **P value** |
|---|---|---|
| Interaction | F (1, 30) = 1,262 | P=0,2701 |
| Treatment | F (1, 30) = 19,34 | P=0,0001 |
| Genotype | F (1, 30) = 42,17 | P<0,0001 |

| **Two-way ANOVA ELN content** | **F (DFn, DFd)** | **P value** |
|---|---|---|
| Interaction | F (1, 30) = 0,05941 | P=0,8091 |
| Treatment | F (1, 30) = 0,5833 | P=0,4510 |
| Genotype | F (1, 30) = 43,72 | P<0,0001 |

| **Two-way ANOVA Number VSMC** | **F (DFn, DFd)** | **P value** |
|---|---|---|
| Interaction | F (1, 20) = 0,1171 | P=0,7358 |
| Treatment | F (1, 20) = 9,485 | P=0,0059 |
| Genotype | F (1, 20) = 53,93 | P<0,0001 |

| **Two-way ANOVA Total Collagen** | **F (DFn, DFd)** | **P value** |
|---|---|---|
| Interaction | F (1, 45) = 20,81 | P<0,0001 |
| Treatment | F (1, 45) = 2,773 | P=0,1028 |

| | | |
|---|---|---|
| Genotype | F (1, 45) = 4,878 | P=0,0323 |
| **Tukey's multiple comparisons test** | | **Adjusted P Value** |
| VEHICLE:WT vs. VEHICLE:CD | | P<0,0001 |
| VEHICLE:WT vs. CURVER:CD | | P=0,9769 |
| VEHICLE:CD vs. CURVER:CD | | P=0,0005 |

| **Two-way ANOVA Green Collagen** | **F (DFn, DFd)** | **P value** |
|---|---|---|
| Interaction | F (1, 48) = 19,20 | P<0,0001 |
| Treatment | F (1, 48) = 2,121 | P=0,1518 |
| Genotype | F (1, 48) = 17,77 | P=0,0001 |
| **Tukey's multiple comparisons test** | | **Adjusted P Value** |
| VEHICLE:WT vs. VEHICLE:CD | | P<0,0001 |
| VEHICLE:WT vs. CURVER:CD | | P=0,1822 |
| VEHICLE:CD vs. CURVER:CD | | P=0,0015 |

| **Two-way ANOVA Red Collagen** | **F (DFn, DFd)** | **P value** |
|---|---|---|
| Interaction | F (1, 46) = 8,411 | P=0,0057 |
| Treatment | F (1, 46) = 0,4011 | P=0,5296 |
| Genotype | F (1, 46) = 0,3175 | P=0,5759 |
| **Tukey's multiple comparisons test** | | **Adjusted P Value** |
| VEHICLE:WT vs. VEHICLE:CD | | P=0,2564 |
| VEHICLE:WT vs. CURVER:CD | | P=0,8036 |
| VEHICLE:CD vs. CURVER:CD | | P=0,0853 |

#### Supplemental Table S3: Statistical data of Hearts

| **Three way ANOVA Sex effect** | **F (DFn, DFd)** | **P value** |
|---|---|---|
| Treatment | F (1, 97) = 22,70 | P<0,0001 |
| Genotype | F (1, 97) = 15,48 | P=0,0002 |
| Sex | F (1, 97) = 0,1639 | P=0,6865 |
| Treatment x Genotype | F (1, 97) = 11,97 | P=0,0008 |
| Treatment x Sex | F (1, 97) = 6,940 | P=0,0098 |
| Genotype x Sex | F (1, 97) = 0,07919 | P=0,7790 |
| **Treatment x Genotype x Sex** | **F (1, 97) = 6,555** | **P=0,0120** |
| **Šídák's multiple comparisons test** | **Adjusted P Value** | |
| VEHICLE:WT-MALES vs. VEHICLE:WT-FEMALES | P=0,0777 | |
| VEHICLE:CD-MALES vs. VEHICLE:CD-FEMALES | P>0,9999 | |
| CURVER:WT-MALES vs. CURVER:WT-FEMALES | P=0,1547 | |
| CURVER:CD-MALES vs. CURVER:CD-FEMALES | P>0,9999 | |

| **Two-way ANOVA Heart vs Body weight** | **F (DFn, DFd)** | **P value** |
|---|---|---|
| Interaction | F (1, 101) = 20,76 | P<0,0001 |
| Treatment | F (1, 101) = 15,13 | P=0,0002 |
| Genotype | F (1, 101) = 16,37 | P=0,0001 |
| **Tukey's multiple comparisons test** | **Adjusted P Value** | |
| VEHICLE:WT vs. VEHICLE:CD | P <0,0001 | |
| VEHICLE:WT vs. CURVER:WT | P =0,9640 | |
| VEHICLE:WT vs. CURVER:CD | P =0,9995 | |
| VEHICLE:CD vs. CURVER:CD | P <0,0001 | |

| **Two-way ANOVA Thickness LV** | **F (DFn, DFd)** | **P value** |
|---|---|---|
| Interaction | F (1, 25) = 7,988 | P=0,0091 |
| Treatment | F (1, 25) = 3,832 | P=0,0615 |
| Genotype | F (1, 25) = 14,30 | P=0,0009 |
| **Tukey's multiple comparisons test** | **Adjusted P Value** | |
| VEHICLE:WT vs. VEHICLE:CD | P= 0,0009 | |
| VEHICLE:WT vs. CURVER:WT | P=0,9306 | |
| VEHICLE:WT vs. CURVER:CD | P= 0,5941 | |
| VEHICLE:CD vs. CURVER:CD | P= 0,0099 | |

#### Supplemental Table S4: Statistical data of 3-NT and XOR

| **Two-way ANOVA 3-NT LV** | **F (DFn, DFd)** | **P value** |
|---|---|---|
| Interaction | F (1, 36) = 39,09 | P<0,0001 |
| Treatment | F (1, 36) = 13,10 | P=0,0009 |
| Genotype | F (1, 36) = 29,96 | P<0,0001 |
| **Tukey's multiple comparisons test** | **Adjusted P Value** | |
| VEHICLE:WT vs. VEHICLE:CD | P<0,0001 | |
| VEHICLE:WT vs. CURVER:WT | P=0,2959 | |
| VEHICLE:WT vs. CURVER:CD | P=0,5088 | |
| VEHICLE:CD vs. CURVER:CD | P <0,0001 | |

| **Two-way ANOVA 3-NT Aorta** | **F (DFn, DFd)** | **P value** |
|---|---|---|
| Interaction | F (1, 28) = 12,12 | P=0,0017 |
| Treatment | F (1, 28) = 11,27 | P=0,0023 |
| Genotype | F (1, 28) = 7,793 | P=0,0093 |
| **Tukey's multiple comparisons test** | **Adjusted P Value** | |
| VEHICLE:WT vs. VEHICLE:CD | P=0,0008 | |
| VEHICLE:WT vs. CURVER:WT | P=0,9997 | |
| VEHICLE:WT vs. CURVER:CD | P=0,9756 | |
| VEHICLE:CD vs. CURVER:CD | P =0.0007 | |

| **Two-way ANOVA XOR LV (Inmunofluorescence)** | **F (DFn, DFd)** | **P value** |
|---|---|---|
| Interaction | F (1, 16) = 2,697 | P=0,1201 |
| Treatment | F (1, 16) = 8,725 | P=0,0093 |
| Genotype | F (1, 16) = 8,653 | P=0,0096 |
| **One-way ANOVA XOR LV (Western Blot)** | | |
| Treatment | F (2, 21) = 34,02 | P<0,0001 |
| **Tukey's multiple comparisons test** | **Adjusted P Value** | |
| WT vs. CD | P<0,0001 | |
| WT vs. CD-CURVER | P=0,0145 | |
| CD vs. CD-CURVER | P=0,0001 | |

| **ANOVA table XOR Aorta (Inmunofluorescence)** | **F (DFn, DFd)** | **P value** |
|---|---|---|
| Interaction | F (1, 19) = 5,952 | P=0,0247 |

| | | |
|---|---|---|
| Treatment | F (1, 19) = 11,49 | P=0,0031 |
| Genotype | F (1, 19) = 2,167 | P=0,1574 |
| **Tukey's multiple comparisons test** | **Adjusted P Value** | |
| VEHICLE:WT vs. VEHICLE:CD | P=0,0478 | |
| VEHICLE:WT vs. CURVER:WT | P=0,9002 | |
| VEHICLE:WT vs. CURVER:CD | P=0,5591 | |
| VEHICLE:CD vs. CURVER:CD | P=0,0037 | |
| **One-way ANOVA XOR Aorta (Western Blot)** | | |
| Treatment | F (2, 9) = 22 | P=0,0003 |
| **Tukey's multiple comparisons test** | **Adjusted P Value** | |
| WT vs. CD | P=0,0014 | |
| WT vs. CD-CURVER | P=0,5770 | |
| CD vs. CD-CURVER | P=0,0004 | |

#### Supplemental Table S5: Statistical data of pNRF2

| **One Way ANOVA Heart** | **F (DFn, DFd)** | **P value** |
|---|---|---|
| Treatment | F (2,28) = 8 | P=0.0014 |
| **Tukey's multiple comparisons test** | **Adjusted P Value** | |
| VEHICLE WT vs. VEHICLE CD | P=0.0084 | |
| VEHICLE WT vs. CURVER CD | P=0.8771 | |
| VEHICLE CD vs. CURVER CD | P=0.002 | |

| **Two-way ANOVA Aorta** | **F (DFn, DFd)** | **P value** |
|---|---|---|
| Interaction | F (1, 34) = 19,30 | P=0,0001 |
| Treatment | F (1, 34) = 17,89 | P=0,0002 |
| Genotype | F (1, 34) = 36,27 | P<0,0001 |
| **Tukey’s multiple comparisons test** | **Adjusted P Value** | |
| VEHICLE:WT vs. VEHICLE:CD | P<0,0001 | |
| VEHICLE:WT vs. CURVER:WT | P=0,9995 | |
| VEHICLE:WT vs. CURVER:CD | P=0,5890 | |
| VEHICLE:CD vs. CURVER:CD | P <0,0001 | |

#### Supplemental Table S6: Statistical data of treatment intake and body weight

| Two-way RM ANOVA Intake-MALES | F (DFn, DFd) | P value |
|---|---|---|
| Treatment x Time | F (45, 240) = 0,7963 | P=0,8189 |
| Treatment | F (3, 16) = 0,2570 | P=0,8552 |
| Time | F (6,238, 99,80) = 0,6699 | P=0,6799 |

| Two-way RM ANOVA Intake-FEMALES | F (DFn, DFd) | P value |
|---|---|---|
| Treatment x Time | F (45, 240) = 0,7335 | P=0,8936 |
| Treatment | F (3, 16) = 1,846 | P=0,1794 |
| Time | F (6,061, 96,97) = 0,9496 | P=0,4642 |

| Two-way ANOVA Intake- WT vs CD | F (DFn, DFd) | P value |
|---|---|---|
| Interaction | F (3, 24) = 0,1347 | P=0,9384 |
| Treatment | F (3, 24) = 0,5263 | P=0,6685 |
| Genotype | F (1, 24) = 2,915 | P=0,1007 |

| Two-way ANOVA Body weight- MALES | F (DFn, DFd) | P value |
|---|---|---|
| Interaction | F (3, 102) = 0,9029 | P=0,4425 |
| Treatment | F (3, 102) = 1,219 | P=0,3068 |
| Genotype | F (1, 102) = 92,76 | P<0,0001 |

| Two-way ANOVA Body weight- FEMALES | F (DFn, DFd) | P value |
|---|---|---|
| Interaction | F (3, 62) = 1,681 | P=0,1803 |
| Treatment | F (3, 62) = 1,483 | P=0,2279 |
| Genotype | F (1, 62) = 38,57 | P<0,0001 |

**Figure S1**

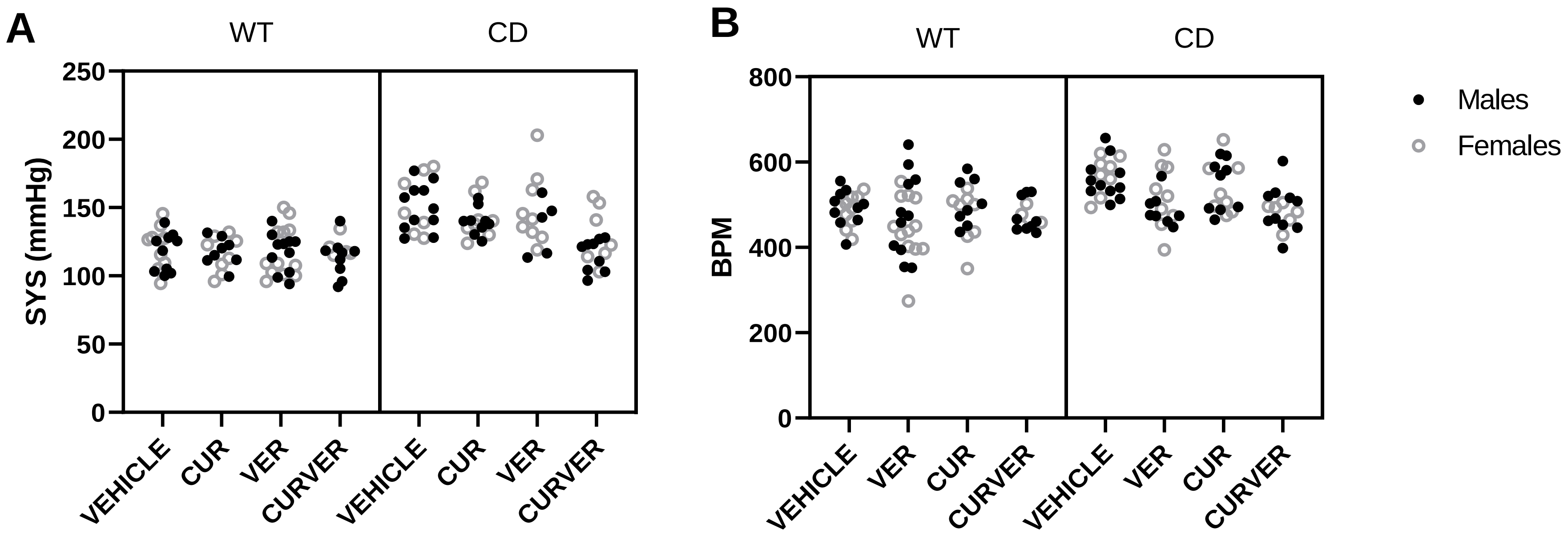
A
WT
CD
250
200
150
100
50
0
SYS (mmHg)
VEHICLE
CUR
VER
CURVER
VEHICLE
CUR
VER
CURVER
B
WT
CD
800
600
400
200
0
BPM
VEHICLE
VER
CUR
CURVER
VEHICLE
VER
CUR
CURVER
Males
Females

Figure S2

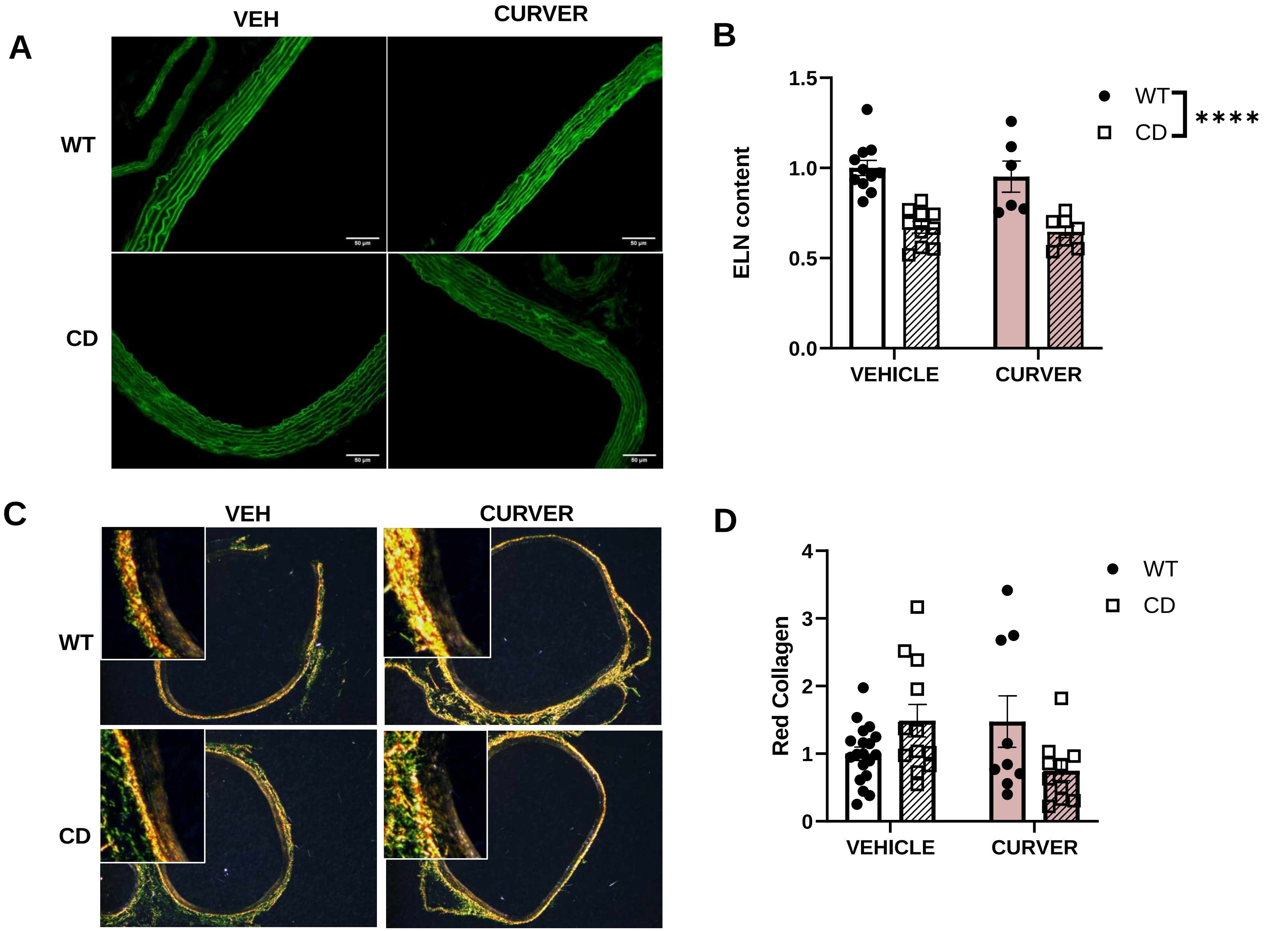
A
VEH
CURVER
WT
CD
B
ELN content
1.5
1.0
0.5
0.0
VEHICLE
CURVER
WT
CD
****
C
VEH
CURVER
WT
CD
D
Red Collagen
4
3
2
1
0
VEHICLE
CURVER
WT
CD

**Figure S3**

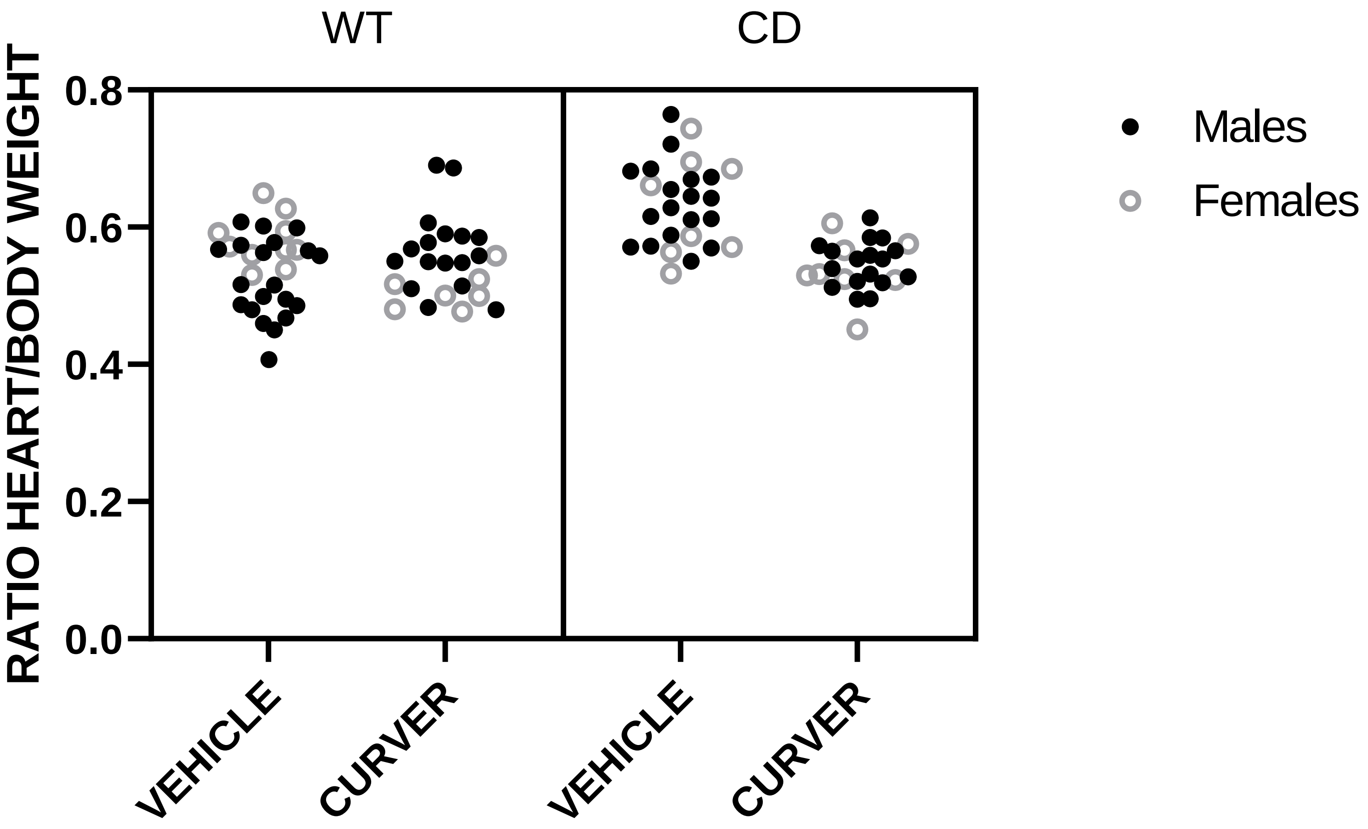

WT
CD
RATIO HEART/BODY WEIGHT
0.8
0.6
0.4
0.2
0.0
VEHICLE
CURVER
VEHICLE
CURVER
Males
Females

### Figure S4

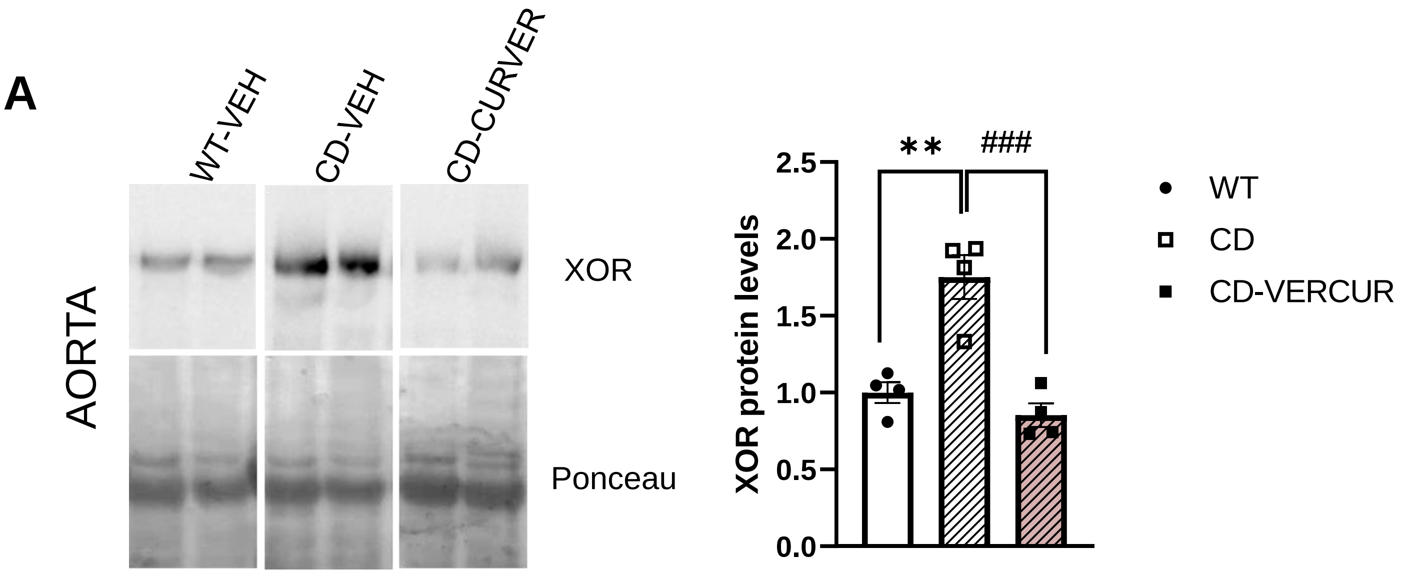
A
WT-VEH
CD-VEH
CD-CURVER
AORTA
XOR
Ponceau
XOR protein levels
**
###
2.5
2.0
1.5
1.0
0.5
0.0
WT
CD
CD-VERCUR

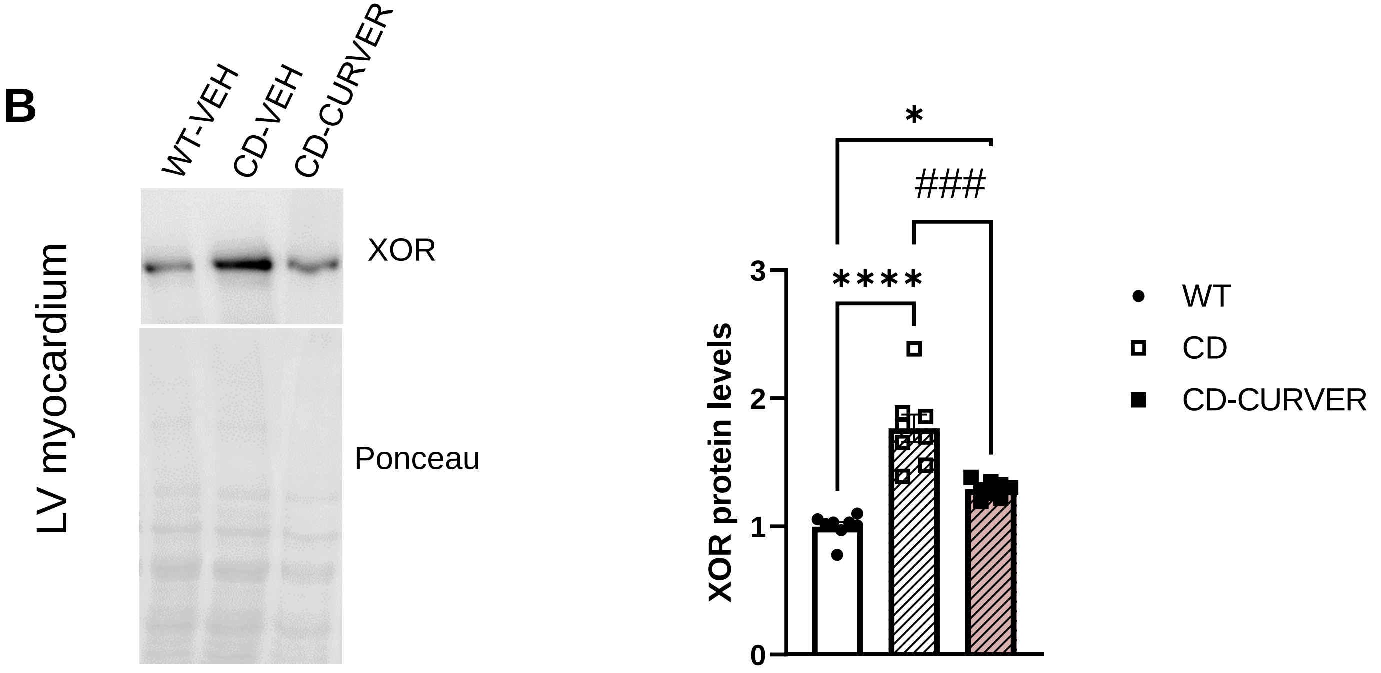
B
WT-VEH
CD-VEH
CD-CURVER
LV myocardium
XOR
Ponceau
XOR protein levels
*
###
****
3
2
1
0
WT
CD
CD-CURVER

Figure S5

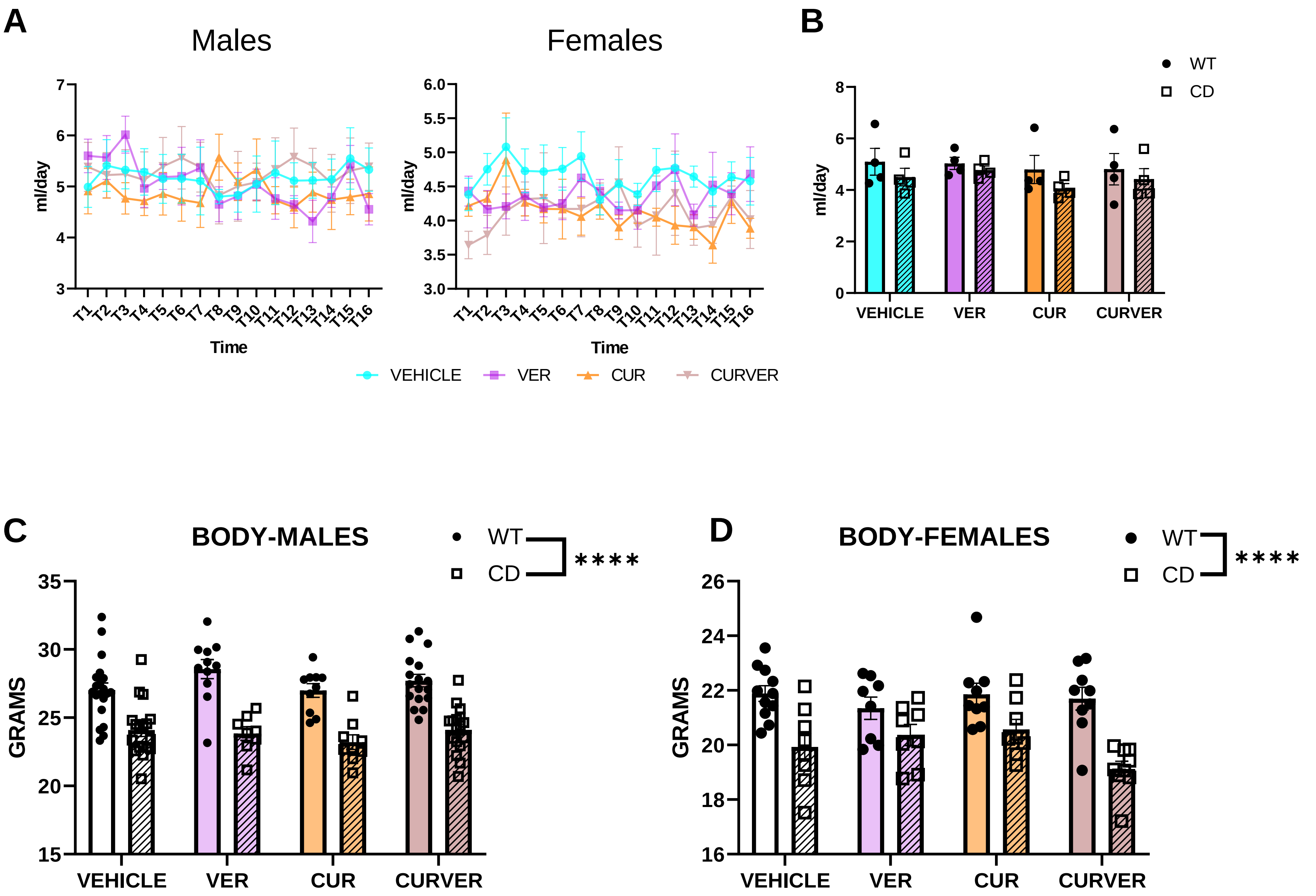

A
Males
Females
7
6
5
4
3
ml/day
6.0
5.5
5.0
4.5
4.0
3.5
3.0
T1 T2 T3 T4 T5 T6 T7 T8 T9 T10 T11 T12 T13 T14 T15 T16
Time
VEHICLE
VER
CUR
CURVER
B
WT
CD
8
6
4
2
0
VEHICLE
VER
CUR
CURVER
C
BODY-MALES
****
GRAMS
35
30
25
20
15
D
BODY-FEMALES
26
24
22
20
18
16

#### Supplemental Figure Legend

##### Figure S1: No effect of sex in SYS and BPM

**(A)** SYS blood pressure and **(B)** BPM measurements in the conscious state using the indirect tail-cuff method in WT (n=14-23) and CD (n=14-18) mice treated with single (CUR, VER) or combined (CURVER) treatments. No significant differences were observed due to sex. Data are represented as mean ± SEM. Three-way ANOVA. Statistical data in Tables S1

##### Figure S2: Analysis of ascending aortas

**(A)** Representative images of green autofluorescence ELN of ascending aortas **(B)** No significant differences were observed in ELN content due to treatment. **(C)** Representative images of ascending aortas under polarized light **(D)** No significant differences were observed in mature (red) collagen content. Data are represented as mean ± SEM. Two-way ANOVA. Statistical data in Tables S2

##### Figure S3: No effect of sex in the ratio heart vs body weight

Data are represented as mean ± SEM. Three-way ANOVA. Statistical data in Table S3

##### Figure S4: Western blot analysis of XOR levels in ascending aortas and LV-myocardium

Representative images of analysed western and the corresponding histograms quantification. Significant difference among groups are showed after one-way ANOVA followed by Tukey's multiple comparisons test for ascending aorta ($F_{(2,9)}$ =22, $p$=0.0003) **(A)** and LV-myocardium ($F_{(2,21)}$ =34.02, $p$<0.0001) **(B)**.

Data are represented as mean ± SEM. * effect of genotype; #, effect of treatment. ***,### p<0.001; ****,#### p< 0.0001. Statistical data in Table S4

##### Figure S5: Treatment intake and body weight

**(A)** The amount of drink per cage was quantified and normalized to the number of animals per cage (2 to 4) and to the time between each change (48 to 60 hours). Consumption was not significantly different between groups ($F_{45,240}$ = 0.7963, $p$ = 0.8184 in males; $F_{45,240}$ = 0.7335, $p$ = 0.8936 in females). **(B)** Daily intake was not significantly different among VEH-WT and the rest of the groups ($F_{3,24}$ = 0.134, $p$ = 0.9384). **(C, D).** None of the treatments changed the reduced body weight presented by CD mice compared to WT mice (effect of genotype $F_{1,102}$ = 92.76, $p$<0.0001 in males; $F_{1,62}$ = 38.57, $p$<0.0001).
